# Modified penetrance of coding variants by *cis*-regulatory variation shapes human traits

**DOI:** 10.1101/190397

**Authors:** Stephane E. Castel, Alejandra Cervera, Pejman Mohammadi, François Aguet, Ferran Reverter, Aaron Wolman, Roderic Guigo, Ivan Iossifov, Ana Vasileva, Tuuli Lappalainen

## Abstract

Coding variants represent many of the strongest associations between genotype and phenotype, however they exhibit inter-individual differences in effect, known as variable penetrance. In this work, we study how *cis*-regulatory variation modifies the penetrance of coding variants in their target gene. Using functional genomic and genetic data from GTEx, we observed that in the general population, purifying selection has depleted haplotype combinations that lead to higher penetrance of pathogenic coding variants. Conversely, in cancer and autism patients, we observed an enrichment of haplotype combinations that lead to higher penetrance of pathogenic coding variants in disease implicated genes, which provides direct evidence that regulatory haplotype configuration of causal coding variants affects disease risk. Finally, we experimentally demonstrated that a regulatory variant can modify the penetrance of a coding variant by introducing a Mendelian SNP using CRISPR/Cas9 on distinct expression haplotypes and using the transcriptome as a phenotypic readout. Our results demonstrate that joint effects of regulatory and coding variants are an important part of the genetic architecture of human traits, and contribute to modified penetrance of disease-causing variants.

## Introduction

Variable penetrance and variable expressivity are common phenomena that cause individuals carrying the same variant to often display highly variable symptoms, even in the case of Mendelian and other severe diseases driven by rare variants with strong effects on phenotype (Chen et al., 2016). For our purposes, we use the term variable penetrance as a joint description of both variable expressivity (severity of phenotype) and penetrance (proportion of carriers with phenotype). These phenomena are a key challenge for understanding how genetic variants manifest in human traits, and a major practical caveat for the prognosis of an individual’s disease outcomes based on their genetic data. However, the causes and mechanisms of variable penetrance are poorly understood. In addition to environmental modifiers of genetic effects, a potential cause of variable penetrance involves other genetic variants with additive or epistatic modifier effects (Cooper et al., 2013). While some studies have successfully mapped genetic modifiers of, for example, BRCA variants in breast cancer (Milne and Antoniou, 2011) and RETT variants in Hirschsprung’s disease (Emison et al., 2005), genome-wide analysis of pairwise interactions between variants has proven to be challenging in humans. In part, this is because exhaustive pairwise testing of genome-wide interactions typically lacks power and is easily affected by confounders (Wei et al., 2014), and a targeted analysis of a specific variant or gene that is strongly implicated in rare disease typically suffers from a low number of carriers. However, emerging large data sets with functional genomic and genetic data from disease cohorts now enable the genome-wide study of mechanistically justified hypotheses of how combinations of genetic variants may have joint effects on disease risk.

In this study, we analyze how regulatory variants in *cis* may modify the penetrance of coding variants in their target genes via the joint effects of these variants on the final dosage of functional gene product, depending on their haplotype combination (Figs. 1, S1). This phenomenon has been demonstrated to affect penetrance of disease-predisposing variants in individual loci (Alberobello et al., 2011; Amin et al., 2012; Butt et al., 2003; Snozek et al., 2009), and explored in early functional genomic datasets (Dimas et al., 2008; Lappalainen et al., 2011). However, genome-wide evidence of regulatory modifiers of disease risk driven by coding variants has been lacking, alongside a generally applicable theoretical framework and analytical methods to study this phenomenon. In this work, we use population-scale functional genomics and disease cohort data sets to show that genetic regulatory modifiers of pathogenic coding variants affect disease risk. Furthermore, we use genome editing with CRISPR/Cas9 to demonstrate an experimental approach for studying the role of regulatory variants as modifiers of coding variant penetrance. We focus on rare pathogenic coding variants from exome and genome sequencing data that provide the best characterized group of variants with strong phenotypic effects, and common regulatory variants affecting gene expression or splicing. Thus, our analysis integrates these traditionally separate fields of human genetics by considering joint effects that different types of mutations have on gene function.

**Figure 1.**
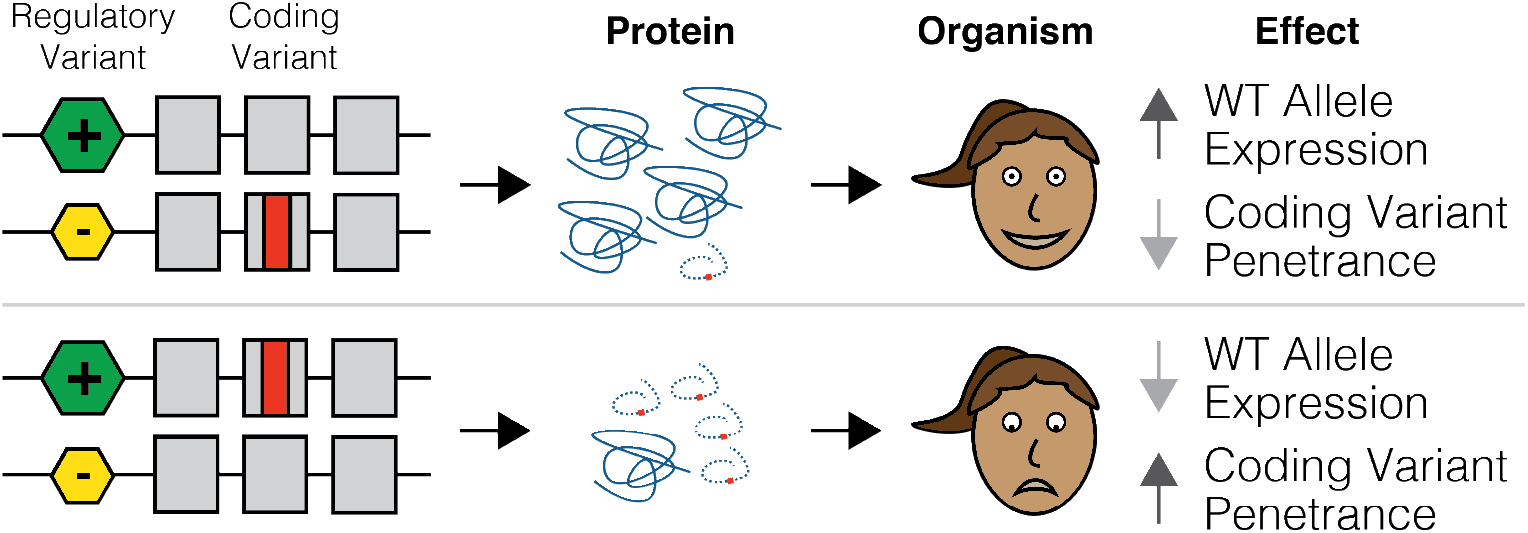
Regulatory variants as modifiers of coding variant penetrance. The hypothesis of this study is illustrated with an example where an individual is heterozygous for both a regulatory variant and a pathogenic coding variant. The two possible haplotype configurations would result in either decreased penetrance of the coding variant if it was on the lower expressed haplotype, or increased penetrance of the coding variant if it was on the higher expressed haplotype. See Figure S1 for a quantitative description of the model.

## Results

### Purifying selection has depleted penetrance increasing haplotype combinations of regulatory and coding variants in the general population

First, we tested the hypothesis that purifying selection should deplete haplotype combinations that increase the penetrance of pathogenic coding variants from the general population. To accomplish this, we analyzed data from the Genotype Tissue Expression (GTEx) project which is representative of the general population in that it lacks individuals with severe genetic disease (GTEx Consortium, 2013). This consists of genotype and RNA-sequencing data of 7,051 samples across 44 tissues from the 449 individuals with exome sequencing and SNP array data of the GTEx v6p release (GTEx Consortium, 2017). Throughout this study, we defined the predicted pathogenicity of variants using their CADD score (see STAR Methods) (Kircher et al., 2014).

We first measured the regulatory haplotype of coding variants using allelic expression (AE) data, which captures *cis* effects of both expression and splice regulatory variation at the individual level (Castel et al., 2015). In our model, purifying selection should result in a depletion of pathogenic variants on higher expressed (Fig. 2a) or exon including (Fig. 2b) haplotypes. For each of the 44 GTEx tissues we calculated the expression of coding variant minor alleles using allelic fold change (aFC) (Mohammadi et al., 2017), and compared the expression of missense variants to allele frequency (AF) matched synonymous controls. Supporting our hypothesis, the minor alleles of missense variants showed reduced allelic expression that was proportional to their predicted pathogenicity (Fig. S2c). Across tissues, rare (AF < 1%) pathogenic (CADD > 15) missense variants showed a significant (p = 4.57e-9) 0.70% reduction of allelic expression compared to synonymous controls, but rare benign (CADD < 15) missense variants did not (p = 0.388) (Figs. 2c, S2a-b). This suggests that in the general population, pathogenic variants are depleted from higher expressed or exon including regulatory haplotypes.

**Figure 2.**
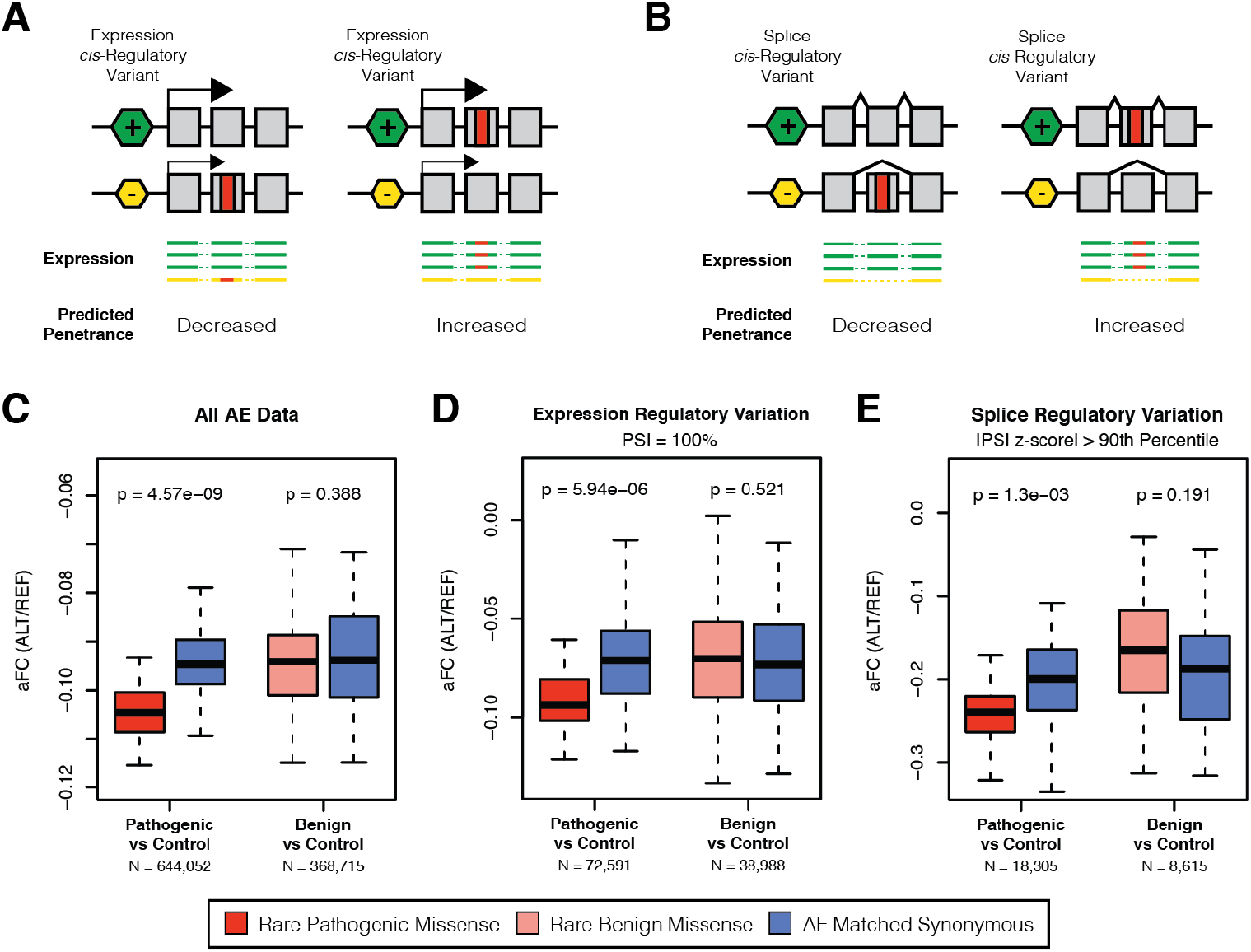
Analysis of regulatory effects at the individual level reveals that pathogenic coding variants are depleted from higher expressed and exon including regulatory haplotypes in the general population. Allelic expression (AE) data can be used to measure the expression of coding variant alleles relative to one another in heterozygous individuals. Reduced expression of the minor allele indicates that the allele is on the lower expressed (A) or spliced (B) haplotype in that individual, potentially reducing its penetrance. C) Boxplot of mean allelic fold change (aFC, log2(alternative allele reads / reference allele reads)) across each of the 44 GTEx tissues calculated for rare (AF < 1%) pathogenic missense (CADD > 15), rare benign missense (CADD < 15), and allele frequency matched synonymous controls. D) Mean aFC across tissues calculated using only variants found in exons where the sample has 100% exon inclusion, as measured by percent spliced in (PSI), which removes allelic effects arising from splice regulatory variation. E) Mean aFC across tissues calculated using only variants found in exons where the sample has substantial variation in exon inclusion compared to the population, as defined by I PS I z-scorel > 90^th^ percentile across all exons, which enriches for allelic effects caused by splice regulatory variation. For each test, the total number (N) of variant aFC measurements across all tissues for pathogenic and benign variants is indicated. P-values are generated by comparing mean aFC of missense variants versus AF matched synonymous controls across tissues using a paired Wilcoxon signed rank test. Outliers are not plotted for ease of viewing. See Figure S2 for tissue level allelic expression results and details of PSI z-score calculation.

In order to study whether this pattern is driven by regulatory variation affecting expression or splicing, which both manifest in allelic expression, we partitioned the coding variants into two groups. To accomplish this, we quantified exon inclusion in each GTEx sample using RNA-seq reads spanning exon junctions to produce a measure of percent spliced in (PSI) for each exon in each sample (Irimia et al., 2014). To isolate the effects of regulatory variation, we analyzed allelic expression only for variants that were found in an exon with 100% inclusion in that individual. As before, rare pathogenic missense variants had significantly reduced expression as compared to synonymous controls (p = 5.94e-6; 1.56% reduction), but rare benign variants did not (p = 0.521), suggesting that pathogenic variants are less likely to accumulate on higher expressed regulatory haplotypes (Fig. 2d). To isolate the effects of splice regulatory variation, we analyzed allelic expression of variants in exons where the sample had substantial deviation in exon inclusion from the population mean. To define these exons, for each exon, a population normalized PSI z-score was produced for each sample allowing for exon inclusion at the sample level to be compared to others (Fig. S2d). When measuring allelic expression of variants found in the top 10% of sample exons by absolute PSI z-score, we again observed that rare pathogenic missense variants had significantly reduced expression as compared to synonymous controls (p = 1.3e-3; 2.00% reduction), but rare benign variants did not (p = 0.191). This suggests that pathogenic variants are less likely to accumulate on haplotypes where the corresponding exon is more likely to be included in transcripts (Fig. 2e). Altogether, these analysis of allelic expression data suggests that in a cohort representative of the general population, pathogenic coding variants less frequently exist in high-penetrance regulatory haplotype combinations, as would be expected under our model.

While allelic expression paired with splice quantification provides a powerful functional readout of latent regulatory variants acting on a gene in each individual, the phenomenon of modified penetrance can also be studied from genetic data alone by analyzing phased haplotypes of coding variants and regulatory variants identified by expression quantitative trait locus (eQTL) mapping in *cis*. Our hypothesis is that in pathogenic coding variant heterozygotes, eQTL-mediated lower expression of the haplotype carrying the “wildtype”, major coding allele increases the penetrance of the rare allele, and vice versa (Figs. 3a, S1). To study this, we developed a test for regulatory modifiers of penetrance that uses phased genetic data (see STAR Methods). Briefly, for each rare coding variant heterozygote, we test whether the major coding allele is on the lower expressed eQTL haplotype (Fig. S3a) and determine if this occurs more or less frequently than would be expected under the null based on eQTL frequencies in the population studied (Fig. S3b). Using simulated data, we found that our test was well calibrated under the null while still being sensitive to changes in haplotype configuration (Fig. S3c-d).

**Figure 3.**
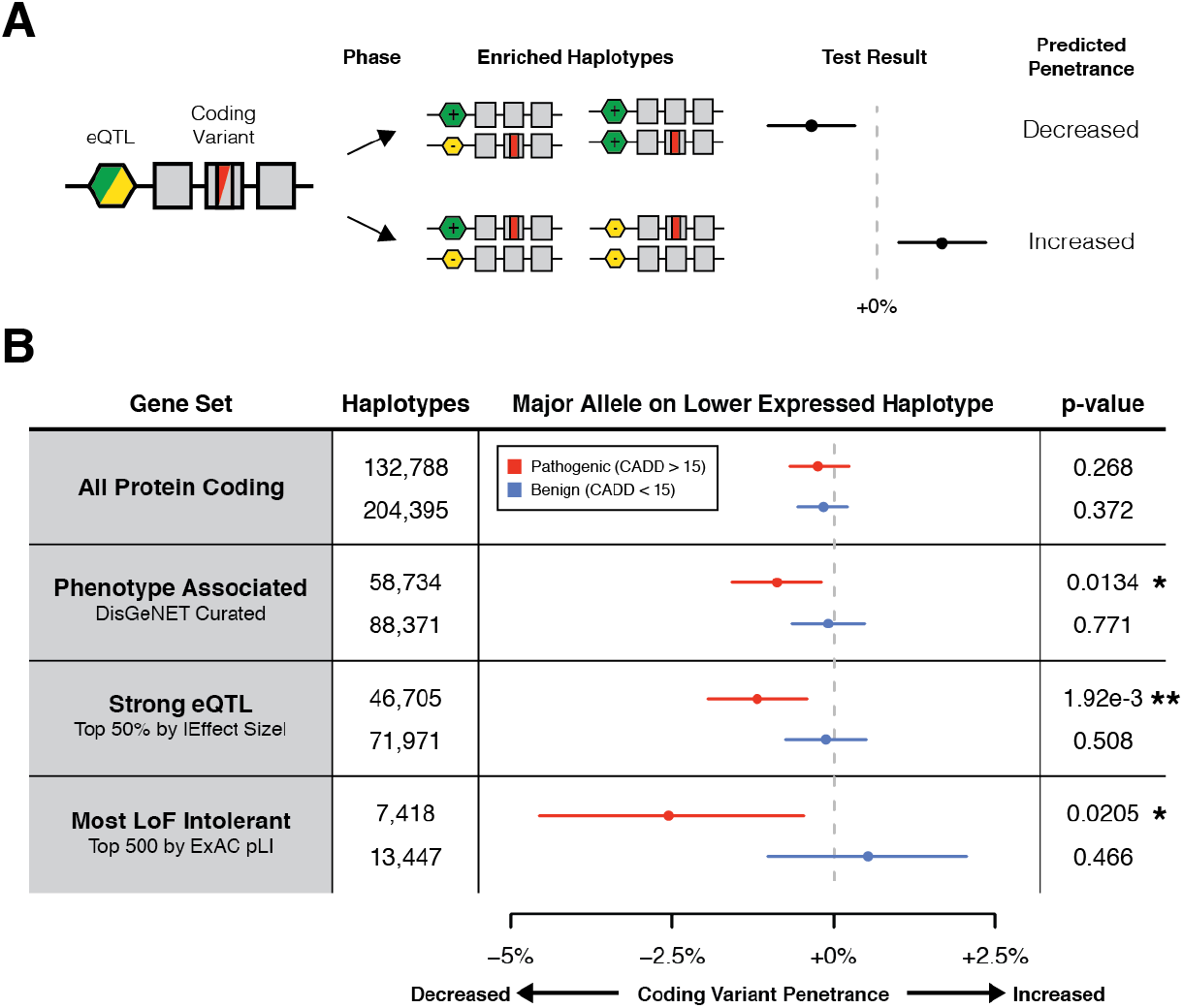
eQTL haplotype configurations that are predicted to increase pathogenic coding variant penetrance are depleted in the genomes of GTEx individuals. **A)** Phased genetic data can be used to produce haplotype configurations between regulatory variation identified using expression quantitative trait locus (eQTL) mapping and coding variant heterozygotes. In our model of modified penetrance, decreased expression of major coding alleles mediated by an eQTL could result in increased penetrance of the minor coding allele and vice-versa (Fig. S1). The frequency of the four classes of eQTL coding variant haplotype configurations are tested against a null distribution, which accounts for eQTL frequencies and assumes that coding variants occur on random haplotypes and in random individuals. This produces a measure of how much more often the major alleles of coding variants are found on lower expressed eQTL haplotypes compared to the null, where a positive value suggests increased penetrance and a negative value decreased penetrance. **B)** Test for regulatory modifiers of coding variant penetrance using 620 GTEx v7 population and read-backed phased whole genomes and GTEx v6p eQTLs, applied to rare (MAF < 1%) pathogenic (CADD > 15, including missense, splice, and stop gained), and rare benign (CADD < 15, including synonymous and missense) SNPs for different gene sets. 95% confidence intervals and empirical p-values were generated using 100,000 bootstraps. * p < 0.05, ** p < 0.01. See STAR Methods and Figure S3 for description and benchmarking of the test, and a description of the gene sets used.

To analyze whether the distribution of coding variants on *cis*-eQTL haplotypes in GTEx showed signs of selection against increased penetrance, we produced a large set of haplotype phased genetic data from GTEx v7, where 30x whole genome sequencing of 620 individuals was available. This was obtained from population based phasing paired with read-backed phasing using DNA-seq reads (Delaneau et al., 2013) and RNA-seq reads (Castel et al., 2016) from up to 38 tissues for a single individual. This allowed us to analyze the haplotypes of 221,487 rare (MAF < 1%) coding variants at thousands of genes with known common (MAF > 5%) eQTLs from GTEx v6p (GTEx Consortium, 2017) (Fig. S4a, Table S1). Using our test for regulatory modifiers of penetrance, we did not observe any significant evidence of reduced penetrance of rare potentially pathogenic variants when all protein coding genes were analyzed together (p = 0.268). However, hypothesizing that selection may be acting primarily at genes that are associated to a phenotype, we focused on a broad set of genes with known phenotypic association (Piñero et al., 2017). For rare potentially pathogenic variants at these genes, we observed a significant (p = 0.0134) decrease of 0.88% in the frequency of haplotypes where the major coding allele was on the lower expressed haplotype expressed than would be expected under the null, while no effect was seen for benign variants (p = 0.771) (Fig. 3b). Similarly, we also observed a significant reduction of predicted penetrance of rare potentially pathogenic variants (p = 1.92e-3) but not benign variants (p = 0.508) in genes with a strong eQTL (top 50% of eQTLs by |effect size|). Finally, we observed the strongest effect at the most loss-of-function intolerant genes as measured by ExAC pLI (-2%, p = 0.0205) (Lek et al., 2016; Samocha et al., 2014), while no effect was seen for benign variants (p = 0.466).

Altogether, combined with observations from functional data of allelic expression, these results suggest that joint effects between regulatory and coding variants have shaped human genetic variation in the general population through purifying selection depleting haplotype combinations where cis-regulatory variants increase the penetrance of pathogenic coding variants (Fig. S1). These patterns are highly significant and consistent, although the genome-wide magnitude of their effects is not strong. However, since our results indicate that regulatory modifiers of penetrance affect primarily pathogenic coding variants, stronger *cis*-regulatory variants, and genes with phenotype effects, genome-wide analysis likely ends up diluting a signal that may be strong and phenotypically relevant for a subset of genes and variants.

### Regulatory modifiers of penetrance affect disease risk in disease cohorts

We next sought to investigate whether regulatory modifiers of penetrance affect disease risk in patients. This would manifest as patients having an overrepresentation of regulatory haplotype configurations that increase penetrance of putatively disease-causing coding variants as compared to controls, where an enrichment of low-penetrance combinations is expected. Importantly, our test is calibrated to eQTL allele frequencies separately in case and control individuals, so that it measures only differences in haplotype configurations and not eQTL frequency between the populations. To test this hypothesis, we applied our genetic test for regulatory modifiers of penetrance to two large disease cohorts in cancer and autism. These diseases have a known contribution from rare coding variants in hundreds of disease-implicated genes, as well as large accessible genomic data sets that include data of rare coding variants and common variants genome-wide.

First, we analyzed autism spectrum disorder (ASD) using the Simons Simplex Collection with genetic data from 2,600 simplex families with one child with autism, their parents, and any unaffected siblings (Fischbach and Lord, 2010). We combined exome sequencing data with imputed genome-wide SNP array data, and used transmission phasing to define haplotypes (Iossifov et al., 2014; Sanders et al., 2015). Common regulatory variants were annotated based on the most significant eQTL variant for each gene in GTEx v6p. This produced joint regulatory and coding haplotypes for 2,304 ASD affected probands (Fig. S4d) and 1,712 of their unaffected siblings (Fig. S4c). We compared the haplotype configurations of rare potentially pathogenic variants observed probands versus unaffected siblings. To enrich for disease causing variants, for probands we analyzed only variants that were not observed in unaffected siblings. We found that at 493 ASD implicated genes (see STAR Methods), probands had a significant increase of major coding alleles of rare potentially pathogenic variants found on lower expressed haplotypes as compared to unaffected siblings (+3.55%; p = 5.12e-3; Fig. 4a). No effect was observed for rare potentially pathogenic variants at control genes matched for the rate of coding variant occurrence and frequency (p = 0.814) or rare benign variants at either ASD implicated (p = 0.459) or control genes (p = 0.840) (Fig. 4a). This suggests that common regulatory variants modify the penetrance of rare coding variants underlying autism, with a joint effect on autism risk.

**Figure 4.**
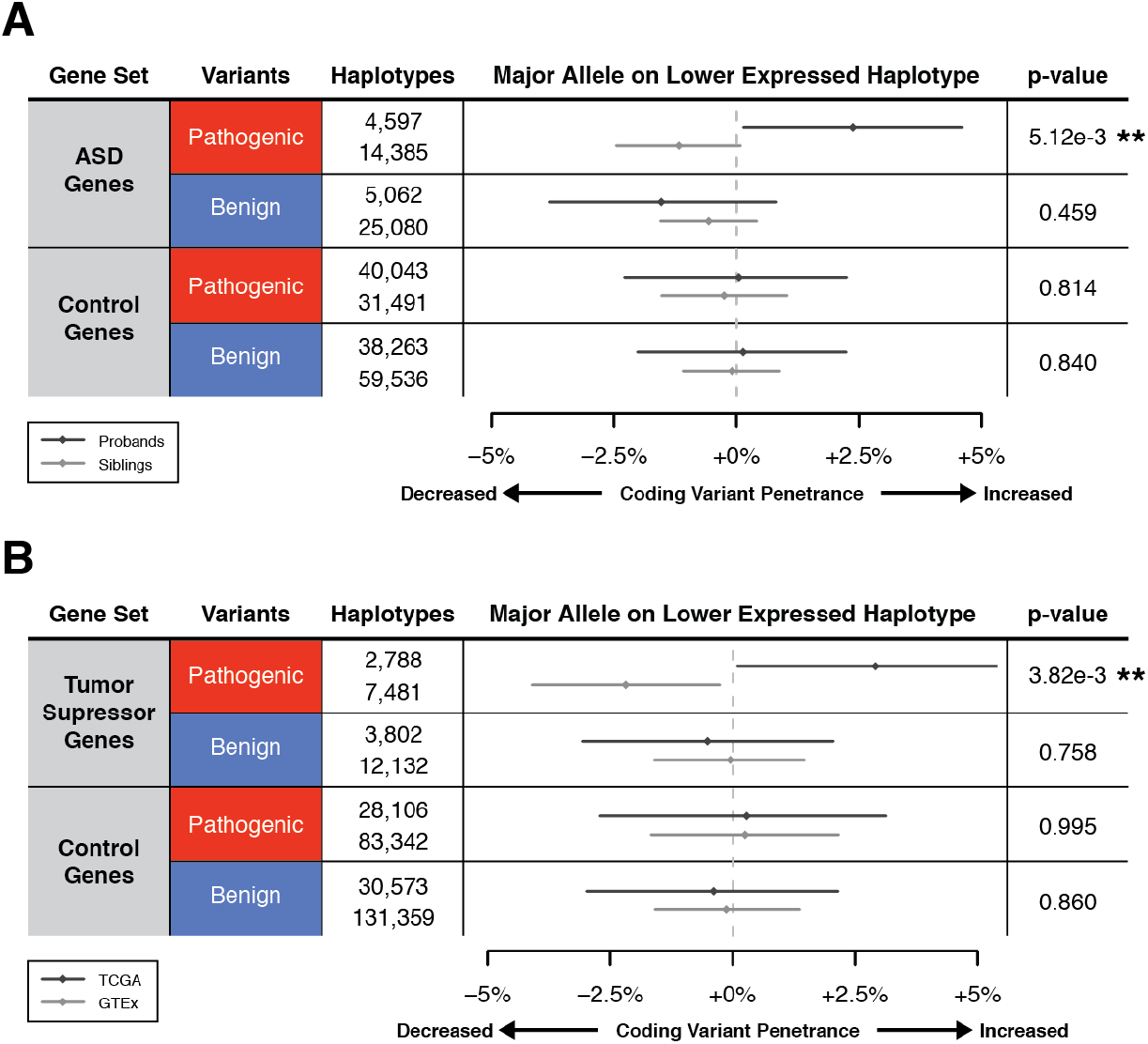
eQTL haplotype configurations that are predicted to increase pathogenic coding variant penetrance are enriched in individuals with disease compared to controls. Comparison of eQTL coding variant haplotype configuration between cases and controls for autism spectrum disorder (ASD) (A) or cancer (B), using the top GTEx v6p eQTL per gene by p-value across all tissues. A) For ASD analysis, haplotype configurations generated from transmission phased genetic data of 2,304 SSC ASD affected probands was compared to haplotype configurations generated from 1,712 of their unaffected siblings. Results for rare (MAF < 1%) pathogenic (CADD > 15, including missense, splice, and stop gained, and frame-shift) and rare benign (CADD < 15, including synonymous and missense) variants are shown for probands and siblings at either ASD implicated or control genes. B) For cancer analysis, haplotype configurations generated from population and read-back phased germline whole genomes of 615 TCGA individuals was compared to haplotype configurations generated from 620 whole genomes of v7 GTEx individuals that are representative of the general population. Results for rare pathogenic (CADD > 15, including missense, splice, and stop gained) and rare benign (CADD < 15, including synonymous and missense) SNPs are shown for TCGA and GTEx at either tumor suppressor genes, or control genes. 95% confidence intervals were generated using 100,000 bootstraps, and empirical p-values for cases versus controls were generated from these confidence intervals (* p < 0.05, ** p < 0.01). Control genes were selected to have within ± 5% the number of coding variants, coding variant frequency, and number of eQTL coding variant haplotypes as disease genes, and had a matched number of haplotypes sampled from them. To enrich for putatively disease-causing variants, for cases, only those variants not observed in controls were used. See STAR Methods for description of gene sets used. See Figure S4 for description of eQTL coding variant haplotypes used for the analysis.

Second, we investigated the role of regulatory modifiers of penetrance in germline cancer risk using genetic data from the Cancer Genome Atlas (TCGA) (Cancer Genome Atlas Research Network et al., 2013). In this analysis, we analyzed tumor suppressor genes that are known to harbor germline risk variants for cancer, often with a dosage-sensitive disease mechanism (Payne and Kemp, 2005). For 615 individuals across 15 cancers where whole genome sequencing reads were available to us (Table S2), we called germline variants and phased these using population (Abecasis et al., 2015) and read-backed phasing (Castel et al., 2016), and analyzed haplotypes of coding variants and common regulatory variants annotated based on the most significant eQTL variant for each gene in GTEx v6p (Fig. S4b). We then compared the haplotype configurations of rare potentially pathogenic variants observed in TCGA individuals versus GTEx samples. To enrich for disease causing variants, for TCGA individuals we analyzed only those variants that were not observed in GTEx samples. We found that at tumor suppressor genes (see STAR Methods), TCGA individuals had a significant increase of major coding alleles of rare potentially pathogenic variants found on lower expressed haplotypes as compared to GTEx controls (+5.10%; p = 3.82e-3; Fig. 4b). No effect was observed for rare potentially pathogenic variants at control genes matched for the rate of coding variant occurrence and frequency (p = 0.995) or rare benign variants at either tumor suppressor genes (p = 0.758) or control genes (p = 0.860). This suggests that increased penetrance of pathogenic germline coding variants by regulatory variation increases cancer risk.

### Experimental demonstration of regulatory modifier effect in the FLCN gene

Our population scale analyses provide observational evidence that regulatory modifiers of penetrance play a role in the genetic architecture of human traits. We next sought to demonstrate an experimental approach for testing this hypothesis for a specific gene by using CRISPR/Cas9 to introduce a coding variant on distinct regulatory haplotypes, followed by quantification of its penetrance from a cellular readout. Our finding that modified penetrance of germline variants by eQTLs may be involved in cancer risk lead us to study a missense SNP (rs199643834, K>R) in the tumor suppressor gene folliculin *(FLCN)* that has a common eQTL in most GTEx v6p tissues (GTEx Consortium, 2017). This SNP causes the Mendelian autosomal dominant disease Birt-Hogg-Dubé Syndrome (Toro et al., 2008) that results in characteristic benign skin tumors, lung cysts, and cancerous kidney tumors and shows variable penetrance (Khoo et al., 2002). We edited the SNP in a fetal embryonic kidney cell line (293T), which is triploid at the *FLCN* gene and harbors a single copy of a common (1000 Genomes AF = 0.428) loss of expression eQTL (rs1708629) located in the 5’ UTR of the gene (GTEx Consortium, 2017; Lin et al., 2014). This variant is among the most significant variants for the *FLCN* eQTL signal, overlaps promoter marks across multiple tissues, and alters motifs of multiple transcription factors (Ward and Kellis, 2012), thus being a strong candidate for the causal regulatory variant of the *FLCN* eQTL (Fig. S5a). We recovered monoclonal cell lines, genotyped them by targeted DNA-seq and performed targeted RNA-seq of the edited SNP (Figs. 5a, S5b). Allelic expression analysis showed that the haplotypes in the cell line are indeed expressed at different levels, likely driven by rs1708629 or another causal variant tagged by it, and the allelic expression patterns allowed phasing of the coding variant with the eQTL (Fig. 5b). In this way, we obtained four clones with a single copy of the Mendelian variant on the lower expressed haplotype (snpLOW), three clones with a single copy on the higher expressed haplotype (snpHIGH), two monoallelic clones with three copies of the alternative allele, and four with only the reference allele (WT) of rs199643834. As a phenotypic readout, we performed RNA-seq on all monoclonal lines.

Using the transcriptomes of these clones, we carried out differential expression analysis. Introduction of the Mendelian SNP had a genome-wide effect on gene expression, with 664 of 20,507 tested genes being significantly (FDR < 10%) differentially expressed in clones monoallelic for the SNP versus wildtype controls (Fig. S5c, Table S3). Gene set enrichment analysis (Wang et al., 2017) of differential expression test results revealed significant (FDR < 10%) enrichment of pathways related to cell cycle control, DNA replication, and metabolism, consistent with the annotation of FLCN as a tumor suppressor, and the occurrence of tumors in patients with the mutation (Table S4). To study the joint effect of the eQTL and Mendelian variant, we quantified the differential expression of these 664 genes in low and high edited SNP expression clones separately (Fig. 5a). As we predicted, clones with higher expression of the SNP showed a significantly stronger differential expression of both downregulated (median = 8.10% increase; 95% CI = 5.93% to 10.36%; p = 8.60e-14) and upregulated (median = 6.52% increase; 95% CI = 4.76% to 8.22%; p = 4.40e-11) genes compared to lower SNP expression clones (Figs. 5c-d). These results provide experimental demonstration that an eQTL can modify the penetrance of a disease-causing coding variant, and suggests a genetic regulatory modifier mechanism as a potential explanation of variable penetrance of rs199643834 in Birt-Hogg-Dubé syndrome.

**Figure 5.**
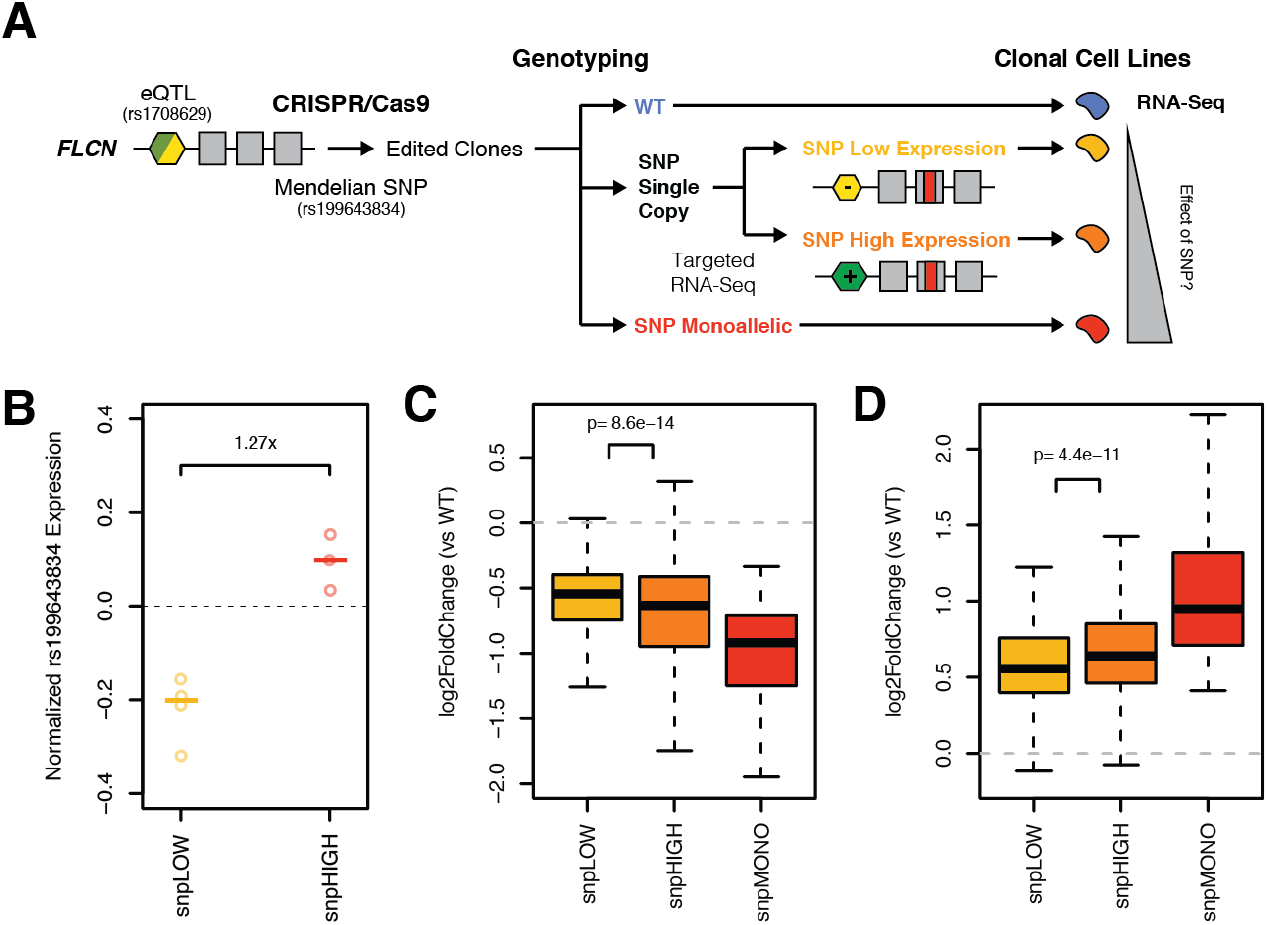
Haplotype aware genome editing of a Mendelian disease SNP in *FLCN* demonstrates that expression regulatory variation can modify its penetrance. **A)** Illustration of the experimental study design, where CRISPR/Cas9 was used to edit a Mendelian missense SNP in *FLCN* (rs199643834) that causes Birt-Hogg-Dubé Syndrome into 293T cells that harbor a single copy loss of expression eQTL for the gene (rs1708629). Monoclonal cell lines were produced, genotyped using targeted DNA-seq of the edit site, and classified as monoallelic for the edit SNP (snpMONO), or as having a single copy. Targeted RNA-seq and allelic expression analysis of the edit SNP was performed for single copy clones, allowing the phase of the SNP with respect to the eQTL to be determined. Using the transcriptome as a phenotype, changes in gene expression compared to wild-type should be stronger in snpHIGH clones versus snpLOW clones if SNP penetrance is modified by the eQTL. **B)** Copy number normalized expression of the edited SNP as measured by targeted RNA-seq (allelic expression, log2(ALT/REF)) in snpLOW (allelic expression < 0, p-value < 0.01, derived from binomial distribution without imbalance) and snpHIGH (allelic expression > 0, p-value < 0.01) clones. **C-D)** Change in expression of genes that were significantly downregulated (**C**, 277 genes) or upregulated (**D**, 387 genes) in clones monoallelic for the edited SNP versus wild-type controls. Single copy edit SNP clones are stratified by haplotype configuration. P-values were calculated using a paired Wilcoxon signed rank test. See Figure S5 for details of the targeted genotyping and RNA-seq, and differential expression analysis results.

## Discussion

In conclusion, we have studied the hypothesis that regulatory variants in *cis* can affect the penetrance of pathogenic coding variants. We used diverse data types from population and disease cohorts, and experimental approaches that together provide strong evidence for our model of modified penetrance due to joint functional effects of regulatory and coding variants. A key component of our work was the integrated analysis of rare coding variants and common regulatory variants, which are too often considered as separate domains in human genetics, despite the fact that their interplay is gaining increasing interest (Werling et al., 2017). Currently, rare coding variants are studied largely by exome sequencing in relatively rare diseases, and common regulatory variant analyses are focused on applications in genome-wide association studies of common diseases. Setting the stage for future studies, our work provides one of the few concrete and generalizable models of modified penetrance of genetic variants in humans, with a clear biological mechanism based on the net effect of variants on the dosage of functional gene product, and is supported by solid empirical analysis of genome-wide genetic data.

Our functional genomic and genetic analysis of the general population provides evidence that purifying selection is acting on joint regulatory and coding variants haplotypes. Importantly, this suggests that the combination of an individual’s regulatory and coding variant genotypes has an effect on phenotype, since purifying selection acts only on traits that affect fitness. Our analyses of autism and cancer cohorts provide direct evidence that regulatory modifiers of coding variants contribute to disease risk, with further indication of regulatory modifiers in the Mendelian Birt-Hogg-Dubé syndrome provided by our experimental approach. The approaches introduced here can be applied to additional disease data sets, with GTEx data providing an essential resource of regulatory variants to empower these analyses. In individual genes, finding regulatory modifiers will require relatively large data sets, and studies of large families with segregating coding variants may be a particularly powerful approach. Genome editing experiments, as we demonstrated for *FLCN*, will be important for functionally validating results from computational analysis.

This work opens additional important areas for future research. Our results demonstrate that the strength of modified penetrance depends on the functional importance and dosage-sensitivity of the gene, effect size of the regulatory variants that affect expression or splicing, and the type of coding variant. Larger data sets are needed to uncover this full spectrum at the level of individual genes instead of gene classes analyzed here. In this work, we focused on loss-of-function analysis, where the expression level of the *non-mutant* haplotype matters, but it is likely that for less common gain-of-function germline and somatic variants, modified penetrance may depend on the expression of the *mutant* haplotype instead. This may be an important consideration for potential future work on variable penetrance of somatic variants in cancer. The dynamics of natural selection on haplotype combinations will be an interesting area of population genetic analysis, where an individual’s fitness depends on multiple variants on different homologs, as well as linkage disequilibrium between these variants.

Finally, we highlight that while other mechanisms are also likely to contribute to variable penetrance of coding variants, analysis of cis-regulatory modifiers is particularly tractable, with multiple practically feasible approaches introduced in this work. Our findings highlight the importance of considering coding variation in the context of regulatory haplotypes in future studies of modified penetrance of genetic variants affecting disease risk.

## Acknowledgements

We would like to thank members of the Lappalainen lab for discussion surrounding the project, and both Kristin Ardlie and Sampsa Hautaniemi who supervised F.A. and A.C. respectively. We thank the GTEx donors for their contributions to science, the GTEx Laboratory, Data Analysis, and Coordinating Center (LDACC), and the GTEx analysis working group (AWG) for their work generating the resource. In particular we would like to thank Ayellet Segre and Xiao Li at the Broad for their work performing WGS variant calling and phasing of GTEx v7 data. The Genotype-Tissue Expression (GTEx) Project was supported by the Common Fund of the Office of the Director of the National Institutes of Health, and by NCI, NHGRI, NHLBI, NIDA, NIMH, and NINDS. We would also like to acknowledge the families at the participating Simons Simplex Collection (SSC) sites, the principal investigators at each site, the coordinators and staff at the SSC sites, the SFARI staff, and the UMACC. Funds for the SSC were provided by the Simons Foundation. Additionally, we would like to acknowledge the contribution of TCGA specimen donors, and The Cancer Genome Atlas Research Network for their analyses. Funds for the TCGA were provided by Cancer Institute and the National Human Genome Research Institute. T.L. and S.E.C. were supported by the NIGMS grant R01GM122924 and NIMH grant R01MH101814, T.L., S.E.C., and P.M. were supported by the NIH contract HHSN2682010000029C, T.L. and P.M. were supported by NIMH grant R01MH106842, and T.L. was supported by the NIH grant UM1HG008901 and 1U24DK112331. A.C. was supported by the Cancer Society of Finland and Academy of Finland grant 284598.

## Author Contributions

S.E.C. and T.L. designed the study and wrote the manuscript. S.E.C., A.V., and T.L. designed analyses and experiments. S.E.C., A.C., F.A., A.W., and A.V. performed analyses and experiments. P.M. aided development of the test for regulatory modifiers of penetrance. F.R. and R.G. provided and assisted in analysis of GTEx PSI data. I.I. provided and assisted in the analysis of SSC data.

## STAR Methods

### Variant Annotation

Variant annotations for SNPs were retrieved from CADD v1.3 (Kircher et al., 2014). As per guidelines by the CADD authors, missense variants with a CADD PHRED score of > 15 were defined as potentially pathogenic. Synonymous variants with a CADD PHRED score < 15 were used as controls. To be considered rare, variants were required to have a MAF < 1% across GTEx v7, 1000 Genomes Phase 3 (1000 Genomes Project Consortium et al., 2015), and gnomAD r2.0.1 (Lek et al., 2016).

### GTEx Allelic Expression Analysis

GTEx v6p allelic expression data generated from whole exome sequencing genotypes were used (GTEx Consortium, 2017). Variants that were in low mapability regions (UCSC mapability track < 1), had less than 10 reads, or had significant (FDR < 1%) evidence that the variant was monoallelic in that individual across all GTEx tissues were excluded (Castel et al., 2015). To minimize the probability that the observed allelic imbalance was due to effects of the AE variants themselves on splicing, only variants farther than 10 bp from an annotated splice site (Kircher et al., 2014) were used. Within each GTEx tissue, when AE measurements from the same variant were present from different individuals, the measurement with the highest read coverage was used. Only variants where the alternative allele was the minor allele were used to ensure that mapping biases were consistent across variants. For missense variants, matched synonymous controls were selected controlling for allele frequency within 25% of missense variants (e.g. between 0.75% and 1.25% for a 1% frequency missense variant).

### GTEx Exon Inclusion Quantification Analysis

Individual level quantifications of exon inclusion were generated for all GTEx v6p samples with the VAST - TOOLS pipeline, which measures the percent spliced in (PSI) of each exon in each individual (Irimia et al., 2014). Within a given tissue, for each exon with at least 10 PSI measurements, PSI z-scores were generated for each sample. Individuals with substantial variation in exon inclusion compared to the population were defined as the top 10% of PSI z-scores across all sample exons (Fig. S2d).

### GTEx Expression Quantitative Trait Loci (eQTL)

The official set of GTEx v6p top significant (FDR < 5%) eQTLs by permutation p-value were used for all analyses such that each gene by tissue had at most a single eQTL (GTEx Consortium, 2017). Those eQTLs where the 95% confidence interval of eQTL effect size overlapped 0, representing weak eQTLs, were discarded (Mohammadi et al., 2017). To produce a single set of cross-tissue top eQTLs, the top eQTL by FDR across tissues was selected for each eGene, with ties broken by choosing the eQTL with the larger effect size. This resulted in a set of 26,942 eGenes each with a single eSNP (Table S1).

### Genetic Data and Haplotype Phasing

GTEx – GTEx v7 genotypes from whole genome sequencing of the 620 individuals who had at least one RNA sample were used. These genomes were population and read-back phased using DNA-seq reads with SHAPEIT2 (Delaneau et al., 2013). Following this, phASER v1.0.0 was used to perform read-backed phasing using RNA-seq reads (Castel et al., 2016) from all samples for each individual, which was a median of 17 tissues, and ranged from 1 to 38. For RNA-seq based read-backed phasing, only uniquely mapping reads (STAR MAPQ 255) with a base quality of ≥ 10 overlapping heterozygous sites were used, and all other phASER settings were left as default. The resulting phased genotypes were imputed into 1000 Genomes Phase 3 (1000 Genomes Project Consortium et al., 2015) with Minimac3 v2.0.1 (Das et al., 2016).

SSC – Genotypes of the SSC cohort from Sanders *et al*. consisting of data generated on Illumina 1 Mv1, 1Mv3, and Omni2.5 arrays (Sanders et al., 2015) were transmission phased using SHAPEIT2 with relatedness data (O’Connell et al., 2014) and then imputed into the 1000 Genomes Phase 3 panel using the Sanger Imputation Service with PWBT (Durbin, 2014; McCarthy et al., 2016). Coding variants called from WES data in lossifov *et al*., were transmission phased on a per variant basis when possible using the genotypes of both parents. The top GTEx eQTLs from across all tissues were used for analysis instead of brain regions only, due to the substantially lower sample sizes in GTEx brain tissues, which result in fewer eQTLs discovered.

TCGA – Paired tumor and normal WGS reads from 925 individuals were used to call germline and somatic variants with Bambino v1.06 (Edmonson et al., 2011). The resulting germline genotypes were population phased with EAGLE2 v2.3 (Loh et al., 2016) using the 1000 Genomes Phase 3 panel (1000 Genomes Project Consortium et al., 2015) and read-back phased with phASER v1.0.0 (Castel et al., 2016). For read-backed phasing, only reads with MAPQ ≥ 30 and with a base quality of ≥ 10 overlapping heterozygous sites were used, and all other phASER settings were left as default. The resulting phased genotypes were imputed into 1000 Genomes Phase 3 (1000 Genomes Project Consortium et al., 2015) with Minimac3 v2.0.1 (Das et al., 2016). Due to the highly variable sequencing depth across TCGA whole genome libraries, from the 925 individuals, 615 individuals with high quality genotyping and phasing were selected for downstream analysis by filtering the bottom 30% of samples by number of variants called and median EAGLE phase confidence across autosomes. This resulted in an approximately equal number of TCGA (615) and GTEx (620) individuals for analyses.

### Test for Regulatory Modifiers of Penetrance Using Phased Genetic Data

Here we test the hypothesis that in loss-of-function coding variant heterozygotes, decreased expression of the major, or “wild type” coding allele mediated by an eQTL can increase the penetrance of the mutant allele by decreasing the dosage of functional gene transcript, and vice-versa (Fig. S1). The null hypothesis is that eQTL mediated changes of major allele expression have no effect on the penetrance of mutant alleles. Since penetrance cannot be easily measured, we instead measure the frequency that the major allele is observed on the lower expressed eQTL haplotype (Fig. S3a). Under the null hypothesis, a coding mutation would occur in random individuals in the population, and on random haplotypes in those individuals, irrespective of their eQTL genotype. Thus, under the null, the frequency of observed major alleles on lower expressed haplotypes would simply be equal to the frequency of the lower expressed eQTL allele in the population. Alternatively, an increased frequency indicates an enrichment of haplotype configurations that increase coding variant penetrance in the population studied, and vice-versa (Fig. S3b). Importantly, the test is calibrated to the eQTL frequency in the specific population studied, so it is internally controlled for differences in, for example, eQTL allele frequencies between cases and controls.

To perform the test, for each observation of a heterozygous coding variant of interest the phased genotypes of the coding variant and the top GTEx cross-tissue eQTL for that gene are used to produce a binary measure of whether the major coding allele is on the lower expressed haplotype (Fig. S3a). Alongside this binary measure the frequency of the lower expressed eQTL allele is recorded.

For each observation of a heterozygous coding variant in a single individual, with genotype *g* let *A* and a denote the higher and lower expressed eQTL alleles, respectively, and *B* and *b* denote the major and minor coding variant alleles, respectively. We assume that the minor allele is the non-functional allele.

For a given haplotype *g*, we define the indicator function *β* such that it is 1 if the functional allele is on a lower expressed eQTL haplotype, and 0 otherwise:

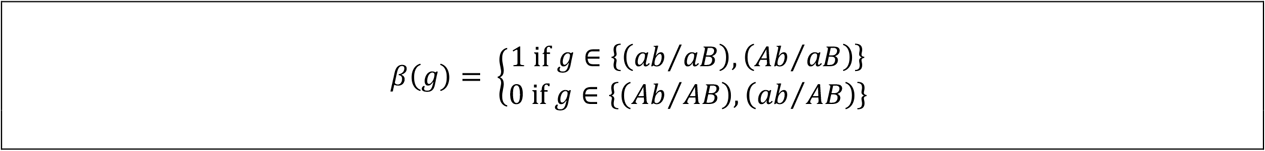

For a given haplotype the expectation for *β* under the null model, where the haplotype configurations are random (H_0_), is:

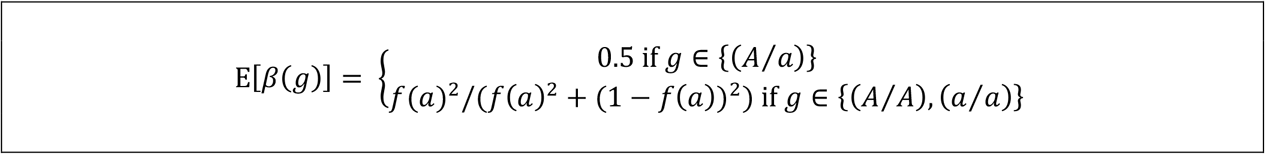

Where *f*(*a*) is the population frequency of the lower expressed eQTL allele included in the tested haplotype *g*.

The indicator function *β* and its expectation under the null model is calculated across all individuals, genes, and variants. The average relative deviation of observed mean of *β* from its expectation was calculated:

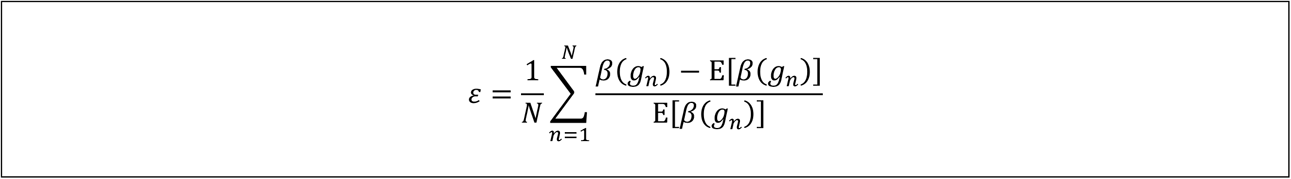

Where *N* is the total number of observed haplotype configurations consisting of an eQTL and coding variant, pooled over all individual, variants, and genes.

Confidence intervals for *ε* are generated by bootstrapping genotypes and the two-sided empirical p-value against H_0_ is calculated as:

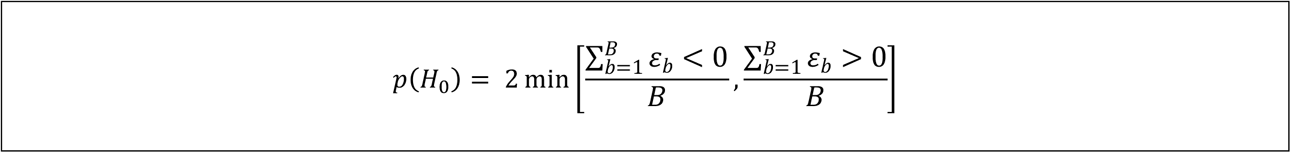

Where *B* is the total number of bootstraps.

We ran the test on simulated haplotype data from 1000 individuals at 500 genes with 1000 replicates. The lower expressed haplotype frequency was set to 50% and the coding variant frequencies as observed in GTEx. This was done across a range of genes exhibiting a bias of major coding alleles being found on lower expressed haplotypes and strengths of this bias. For the test, 1000 bootstrap samples were used. We found that at 5% significance threshold, 5% of simulation replicates were significant, suggesting that the test is well calibrated under the null. For real world data, reported in the study, we used 100,000 bootstrap samples to calculate p-values and derive confidence intervals.

This is a similar problem to that addressed by the Poisson-Binomial distribution, which describes the sum of successes in a set of independent Bernoulli trials with different success rates. However, the bootstrap approach is more convenient for calculating confidence intervals and accounting for differences in sample size between control genes and genes of interest. We compared p-values derived from our test to those derived from a Poisson-Binomial distribution with parameters *E*[*β*(*g*_1_)] … *E*[*β*(*g_N_*)]. In practice, our p-values are very similar to that generated using the Poisson-Binomial distribution (Pearson correlation = 0.996, slope = 0.997, Fig. S3e).

### Gene Sets

A list of phenotype associated genes was produced by downloading all curated gene to phenotype associations from DisGeNET (Piñero et al., 2017) v5.0 on 06/08/17. Genes with strong eQTLs were selected as the top 50% of eGenes by absolute eQTL effect size (Mohammadi et al., 2017). Extremely loss-of-function intolerant genes were selected as the top 500 by ExAC pLI (Lek et al., 2016). A broad set of genes associated with autism spectrum disorder was produced by combining high confidence SFARI database genes (https://gene.sfari.org, categories 1, 2, and S) downloaded on 10/20/17, genes from Krumm *et al*., with nominally significant (p < 0.05) enrichment of *de novo* SNVs in probands versus siblings, and genes with recurrent likely gene disrupting and missense *de novo* mutations in Iossifov *et al*. (Iossifov et al., 2014; Krumm et al., 2015). In total, this resulted in a list of 493 ASD associated genes. A list of 983 down-regulated tumor suppressor genes in tumor samples versus normal tissue in TCGA expression data was downloaded from the Tumor Suppressor Gene Database (Zhao et al., 2015) website (https://bioinfo.uth.edu/TSGene/) on 08/24/17.

### CRISPR/Cas9 Guide Selection and Cloning

Prior to RNA design and editing we verified the genotype at the regions of interest, namely the Mendelian variant rs199643834 and eQTL variant rs1708629. Crude extracts prepared from 293T cells were used to amplify the above regions using forward and reverse genotyping primers FLCN_genot and FLCNeQTL_genot, respectively (Table S5). Amplicons were sequenced by both Sanger sequencing and on the Illumina MiSeq. The 293T cell genotype was Ref/Ref at rs199643834 and Ref/Alt at rs1708629. There were no single nucleotide changes close to rs199643834 that may affect sgRNA activity or require modified homologous template.

Using computational algorithms with prioritization for on-target efficiency and reduced off-target effects (available online: CRISPR Design tool (crispr.mit.edu) and E-CRISPR (Heigwer et al., 2014) we identified Streptococcus pyogenes Cas9 (SpCas9) guide RNAs that bind near variant rs199643834 (A > G). We selected three sgRNA sequences within 50 bp of the target SNP (rs199643834), which were predicted to result in maximum cleavage efficiency without off-target effects (Table S5). Annealed oligomers inclusive of guide RNA sequences were sub-cloned into the lentiCRISPRv2 plasmid (Addgene plasmid #52961), which contains expression cassettes for the guide RNA, a human codon-optimized Cas9, and a puromycin resistance gene (Sanjana et al., 2014). Plasmids were transformed into chemically competent E. coli (One Shot Stbl3 Chemically Competent E. coli, ThermoFisher Scientific, cat#: C737303), and grown at 30°C; plasmid DNA was extracted and purified. A 150 bp single-stranded DNA template (ssODN) for precise editing by homologous recombination (HDR) carrying the rs199643834 A allele was designed and obtained from IDT DNA in the form of lyophilized ultramer (Table S5).

### Transfections and T7 Endonuclease I(T7E1) Assays

Human 293T cell line (ATCC, cat. # CRL-3216) was adapted to and subsequently routinely grown in Opti-MEM/5% CCS (newborn calf serum), 1% GlutaMAX, 1% Penicillin/Streptomycin and sodium pyruvate.

For transfection with Cas9- and sgRNA-expressing plasmids as well as ssODN template, cells were harvested for seeding at a log growth phase (approximately 70% confluency). In a 6-well format, 300,000 293T cells were seeded a day prior to transfection. The next day 2 μg of each lentiCRISPR v2 plasmid and 0.5 μg of ssODN HDR template were delivered into the cells using Lipofectamine 3000 reagent (ThermoFisher Scientific, cat. # L3000008). At 24-hours post-transfection selective pressure in the form of 5 μg/ml puromycin was applied for 8 hours to enrich for transfected cells. The short time-frame reduces the chances of selecting monoclonal lines with stable plasmid integration. Following two days of cell growth cells were harvested and crude extracts prepared from a small fraction for genotyping. The remainder of the cells were frozen for subsequent isolation of cell lines containing desired edits.

For T7E1 assays, a 362 base pair region flanking rs199643834 was PCR-amplified from the crude extracts using FLCN_genot primers and purified using Ampure XP beads (Beckman-Coulter, part #: A63880). Purified products were heteroduplexed, digested with T7 endonuclease 1 (NEB, cat # M0302L), and run on a 2% agarose gel. Cleavage patterns from editing experiments conducted with each sgRNA were qualitatively analyzed to determine each Cas9/sgRNA cutting efficiency to guide further experiments. Subsequently, the crude cell lysates were used to prepare amplicon libraries containing ScriptSeq adapters, which were sequenced on the Illumina MiSeq instrument with paired-end 150 bp reads. Rates of indel mutations by non-homologous end joining (NHEJ) and precise SNP editing by homology-directed repair (HDR) were determined by an in-house analysis pipeline.

### Generation and Identification of Monoclonal Cell Lines Containing Desired Precise Edits

The initial screening showed that editing of 293T polyclonal cell population at rs199643834 with sgRNA 1 resulted in the highest rate of HDR. This population was were selected for single-cell sorting in 96-well format on SONY SH800 to obtain monoclonal edited cell lines. Following 10 days of cell growth, individual wells were scored for the presence of healthy colonies, and altogether approximately 1920 healthy colonies were screened. At first passage a third of the cells from each well were collected for crude cell extracts and genotyping.

High throughput genotyping was performed by preparing an amplicon library from each crude extract with Nextera adapters enabling differential custom dual-indexing. Screening for desired mutations was performed using in-house software. In total, 4 wild-type (Ref/Ref), 7 heterozygous (Ref/Alt) and 2 homozygous mutant (Alt/Alt) clones with each desired mutation were expanded for downstream analyses.

### Targeted RNA-seq of Allelic Series and eQTL Phasing

Expanded lines were grown to 70-80% confluency and RNA was isolated using the Qiagen RNAeasyMini kit. cDNA was synthesized from each RNA sample and the region spanning the Mendelian variant rs199643834 was amplified using primers FLCN_exon9-10-F and FLCN_exon11-R2, containing Nextera adapters (Table S5). Targeted amplicons were dual-indexed using custom Nextera indexes and sequenced on the Illumina MiSeq with 2x150 bp reads.

For all the 13 lines the genotype determined by DNA-sequencing was confirmed by RNA-seq reads. For the 7 lines with a single copy of the edited SNP, we performed allelic expression analysis. Reads were aligned to hg19 using STAR (Dobin et al., 2013). The number of reads mapping to the reference and alternative alleles was quantified using allelecounter requiring MAPQ = 255 and BASEQ ≥ 10 (Castel et al., 2015). Across samples, there was a median of 34,870 reads passing filters overlapping the site. A binomial test using reads containing the edit SNP allele against a null of 1/3 (corresponding to a single copy of the edit SNP) was performed. Copy number normalized allelic expression of the edit SNP was calculated as log2((ALT_COUNT/REF_COUNT)/(1/3)). Samples with allelic expression < 0 and binomial p < 0.01 were categorized as snpLOW (edit SNP on lower expressed eQTL haplotype), and those with allelic expression > 0 and binomial p < 0.01 were categorized as snpHIGH (edit SNP on higher expressed eQTL haplotype).

### RNA-seq and Gene Expression Analysis of Edited 293T Cells

RNA sequencing libraries were prepared using the TruSeq Stranded mRNA Library Sample Preparation Kit in accordance with manufacturer’s instructions. Briefly, 500ng of total RNA was used for purification and fragmentation of mRNA. Purified mRNA underwent first and second strand cDNA synthesis. cDNA was then adenylated, ligated to Illumina sequencing adapters, and amplified by PCR (using 10 cycles). Final libraries were evaluated using fluorescent-based assays including PicoGreen (Life Technologies) and Fragment Analyzer (Advanced Analytics), and were sequenced on the Illumina NovaSeq Sequencing System using 2 x 100bp cycles to a median depth of 52.8 million reads. Trimmomatic (Bolger et al., 2014) v0.36 was used to clip Illumina adaptors and quality trim, and reads were aligned to hg19 using STAR (Dobin et al., 2013) in 2 pass mode. A median of 98% of reads mapped to the human genome, with a median of 95.2% reads mapping uniquely. featureCounts (Liao et al., 2014) v1.5.3 was used in read counting and strand specific mode (-s 2) with primary alignments only to generate gene level read counts with Gencode v19 annotations used in GTEx v6p (GTEx Consortium, 2017). Differential expression analysis was performed using DESeq2 (Love et al., 2014) v1.16.1 and R v3.4.0 on genes with a mean of greater than 5 counts across samples. FDR correction of p-values was performed using Benjamini Hochberg. Gene set enrichment analysis on differential expression data was performed using the Web-based Gene Set Analysis Toolkit (Wang et al., 2017) with Wikipathway enrichment categories.

### Data Use, Availability, and Accessions

GTEx v6p eQTLs are publically available through the GTEx Portal (https://gtexportal.org/). GTEx genotype data, AE data, and RNA-seq reads are available to authorized users through dbGaP (study accession phs000424.v6.p1, phs000424.v7.p2). TCGA data is available to authorized users through dbGap (study accession phs000178.v9.p8). HEK293T RNA-seq data generated in this study is available on the SRA under accession TBD.

## Supplemental Information

### Supplementary Tables

**Table S1**. Top cross tissue GTEx v6p eQTLs per gene.

**Table S2**. TCGA individuals and respective cancer types used for analysis.

**Table S3**. Differentially expressed genes in CRISPR/Cas9 edited rs199643834 monoallelic versus wildtype 293T cells.

**Table S4**. Pathway based gene set enrichment analysis of rs199643834 differential expression data.

**Table S5**. Oligonucleotides used in this study.

### Supplementary Figures

**Figure S1.**
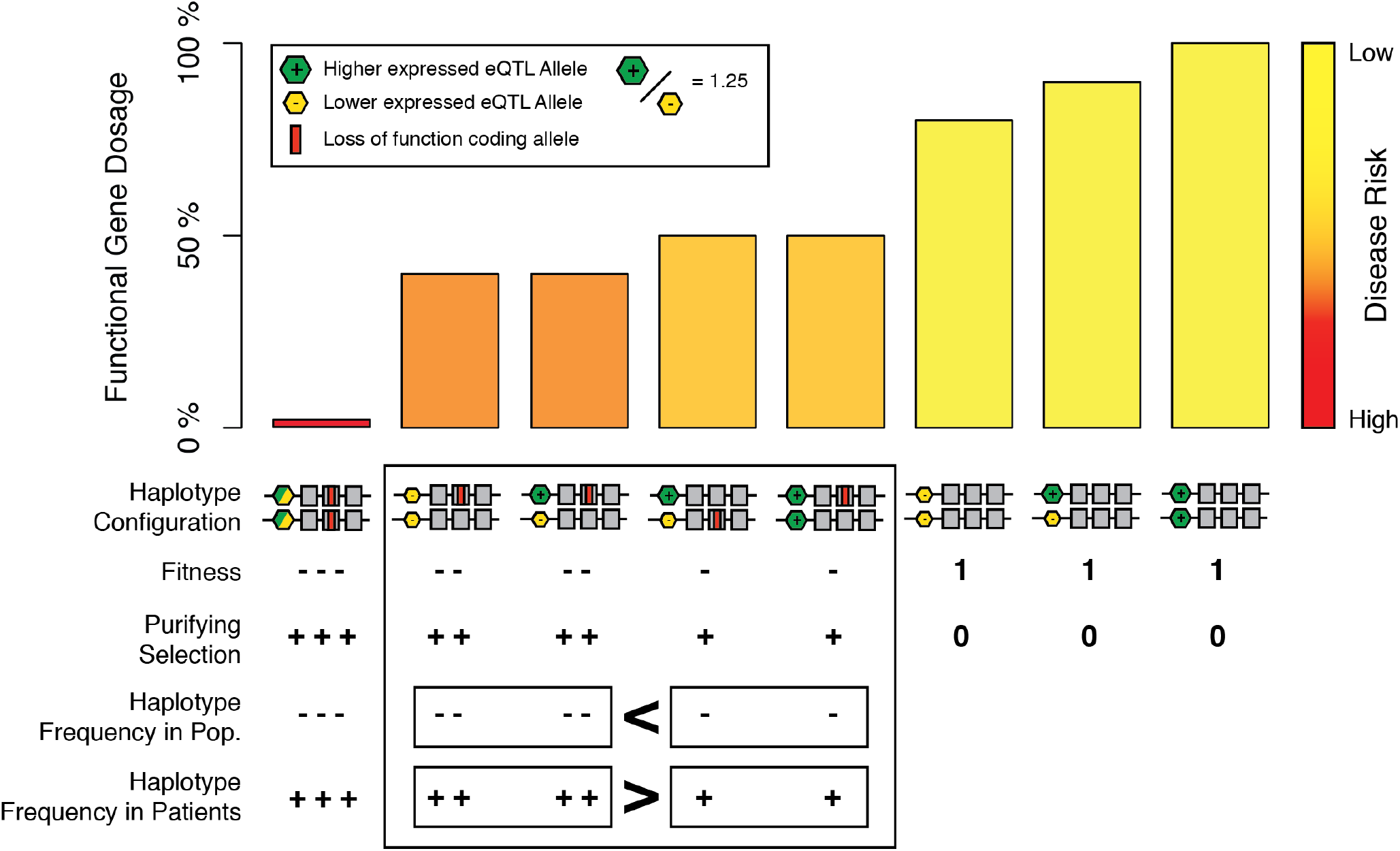
Illustration of the key features of our model of joint effects of regulatory and coding variants on functional gene dosage and selection. Under the model, regulatory variation altering functional gene dosage is particularly important in loss-of-function heterozygotes, where the dosage of functional protein is already reduced to half. Our general assumption is that common regulatory variants typically have such low effects on gene dosage that in the absence of coding variants, they do not cause severe disease or substantial reduction of fitness. Accordingly, in this example, under an additive model of gene expression, the higher expressed eQTL allele increases expression by 1.25x, and disease risk increases non-linearly with decreasing gene dosage, there are potentially large disease risk differences for loss-of-function heterozygotes depending on eQTL haplotype configuration. This results in purifying selection acting more strongly against haplotype configurations that decrease functional gene dosage, while acting more weakly on those that increase functional gene dosage. At the population level this differential strength of purifying selection would result in haplotype configurations that increase functional gene dosage being present at higher frequencies than those that decrease dosage. We note that while we believe that this general model is plausible for many genes with dosage-sensitivity, other scenarios are likely to exist, and for example, fully recessive genes or gain-of-function coding variants would not follow this model. Future work and larger data sets are needed to elucidate the full picture of relative importance of different types of joint effects of regulatory and coding variants.

**Figure S2.**
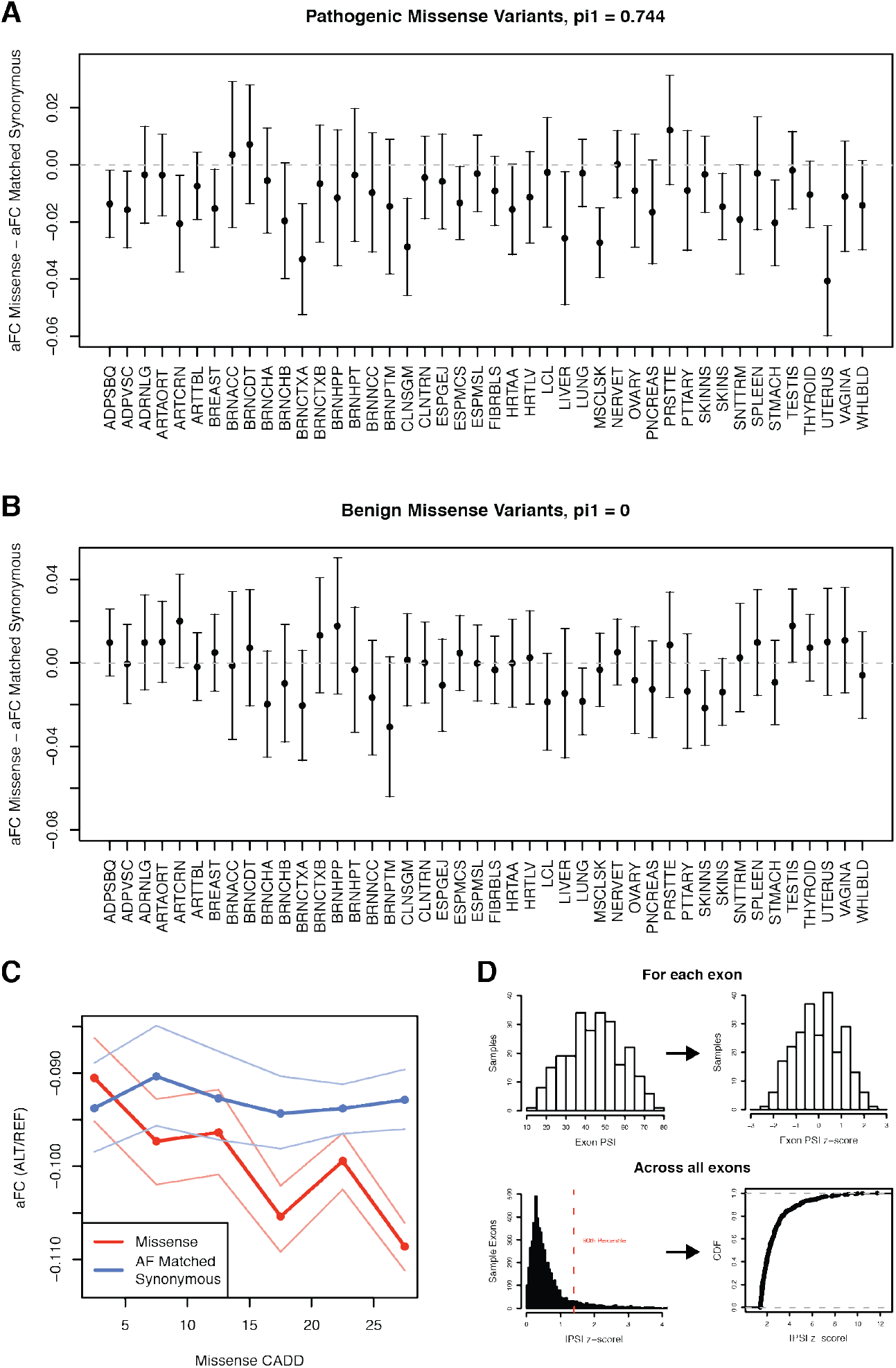
Using GTEx allelic expression (AE) and percent spliced in (PSI) to estimate the penetrance of coding variants at the individual level. Difference in allelic expression between rare (AF < 1%) potentially pathogenic (CADD > 15) **(A)** and benign (CADD < 15) **(B)** missense variants and AF matched synonymous variants across GTEx tissues. A negative difference indicates reduced expression of missense variants compared to synonymous controls. Bars show the 95% confidence interval of the difference of the means between missense and synonymous variants. Storey’s Π_1_ indicates the estimated proportion of true positives across the GTEx tissues. **C)** Allelic expression for rare missense and AF matched synonymous variants in bins of 5 CADD PHRED with 95% confidence interval across GTEx tissues. **D)** Top, illustration of PSI z-score calculation from PSI measurements for the exon HsaEX0054530 in GTEx whole blood. Bottom, histogram of all absolute exon PSI z-scores for coding variants across GTEx whole blood, with 90^th^ percentile shown. Coding variants with an absolute PSI z-score greater than the 90^th^ percentile were considered to be in exons with substantial variation in PSI compared to the population.

**Figure S3.**
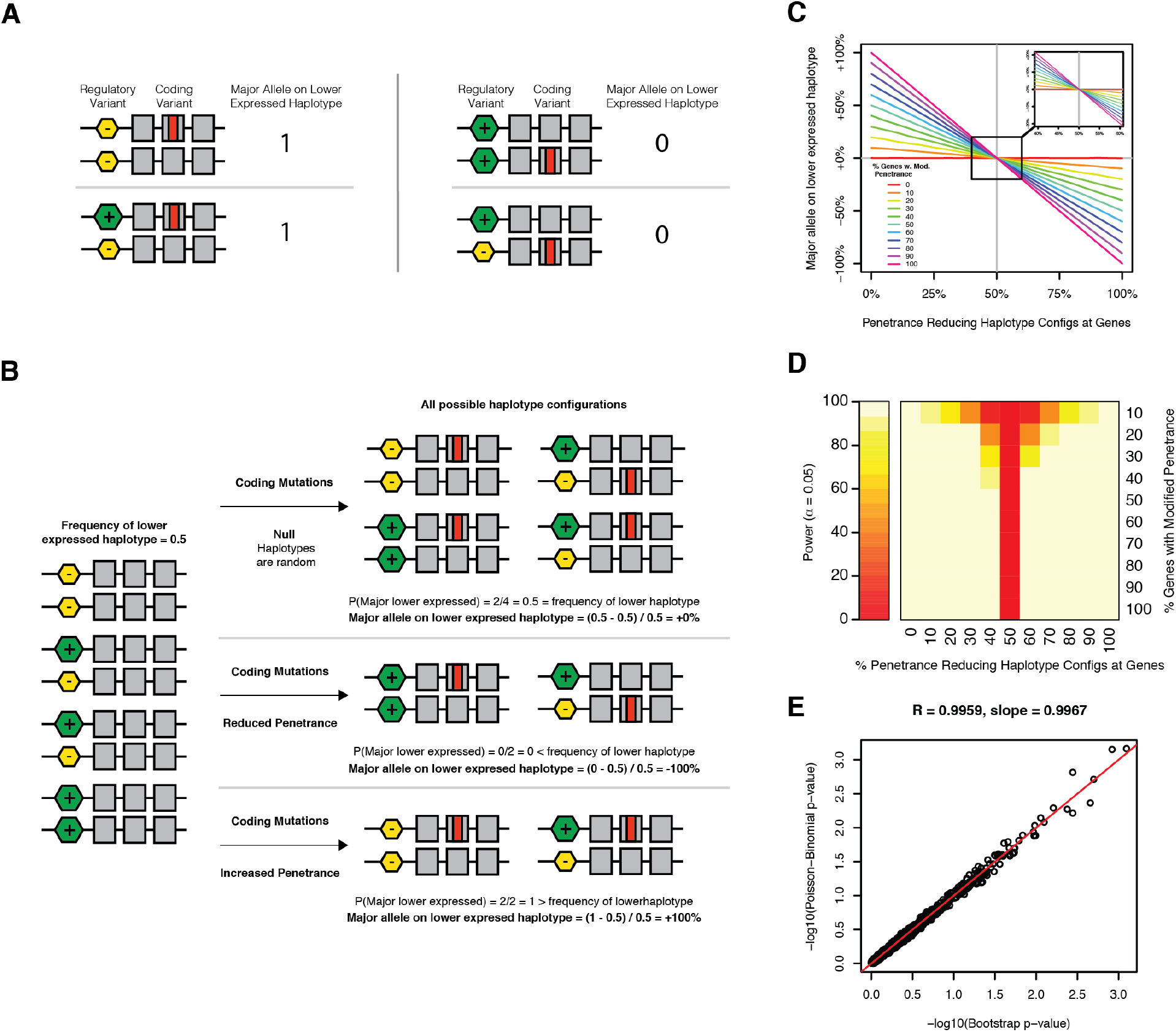
Test for regulatory modifiers of coding variant penetrance using phased genetic data. **A)** As input the test takes phased genotypes of coding variants and the eQTL for that gene. For each individual heterozygous for a coding variant a binary measure is produced to indicate if the major (wild-type) allele is on the lower expressed eQTL haplotype. **B)** Across a population of individuals, the null expectation is that the observed haplotype configurations are a random sampling of all possible configurations, and thus the proportion of observed major alleles on the lower expressed haplotype is equal to the frequency of the lower expressed haplotype in the population. The diagram depicts a single gene example, but observations are aggregated across genes, and the difference between the observed frequency of major alleles on the lower expressed haplotype and the lower expressed haplotype frequency across those genes is calculated. **C)** Results of test performed on simulated haplotype data from 1000 individuals at 500 genes with 1000 replicates using a lower expressed haplotype frequency of 50% and coding variant frequencies observed in GTEx, across a range of genes exhibiting joint effects between regulatory and coding variants and effect size. The simulated effect size is described by the x-axis in terms of the percentage of observed haplotype configurations that decrease penetrance. **D)** Power to detect significant (a = 0.05) regulatory modifiers of penetrance from simulation data in (C) is robust across a range of effect sizes. **E)** Comparison of p-values calculated using either the bootstrap approach or with the Poisson Binomial distribution from 1000 simulations of 1000 haplotypes generated under the null shows that they are extremely similar. The equality line is shown in red. See STAR Methods for more information.

**Figure S4.**
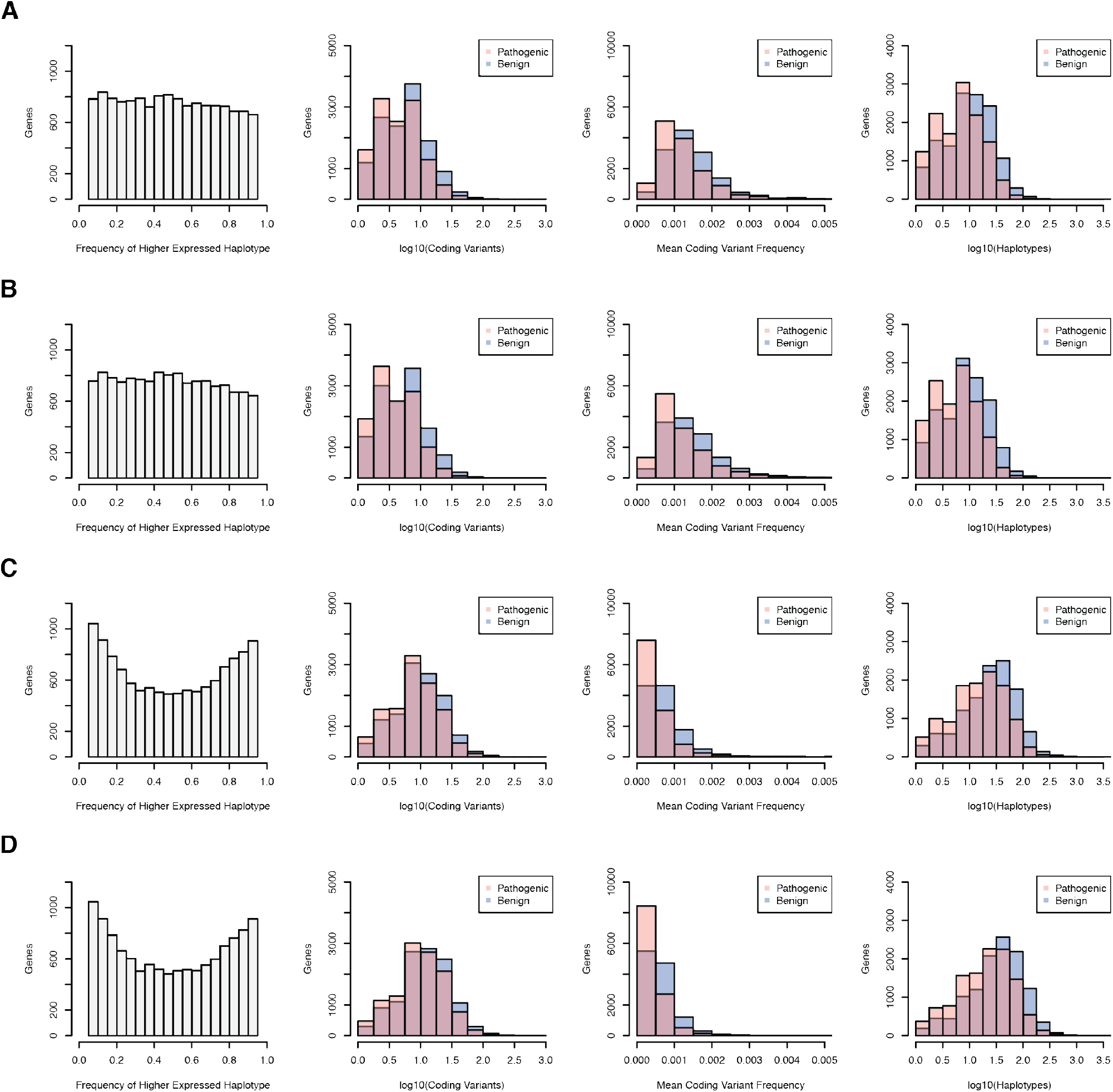
Gene level metrics of common (MAF > 5%) regulatory and rare (MAF < 1%) coding variant haplotypes. Haplotypes generated using potentially pathogenic (CADD > 15) or benign (CADD < 15) rare coding variants and the top cross-tissue GTEx v6p eQTLs to define higher and lower expressed haplotypes. Histograms of higher expressed haplotype frequencies, number of coding variants with haplotype data, mean coding variant frequency, and number of haplotypes observed at the gene level for haplotypes from 620 phased and imputed GTEx v7 whole genomes **(A)**, 615 phased and imputed TCGA germline whole genomes **(B)**, phased and imputed array and whole exome data from 1,712 SSC unaffected siblings **(C)**, and 2,304 SSC probands **(D)**. Differences in the higher expressed haplotype frequency distribution between populations results from differences in eQTL allele frequency. This is not expected to cause systematic bias in our test of modified penetrance shown in Fig. S3.

**Figure S5.**
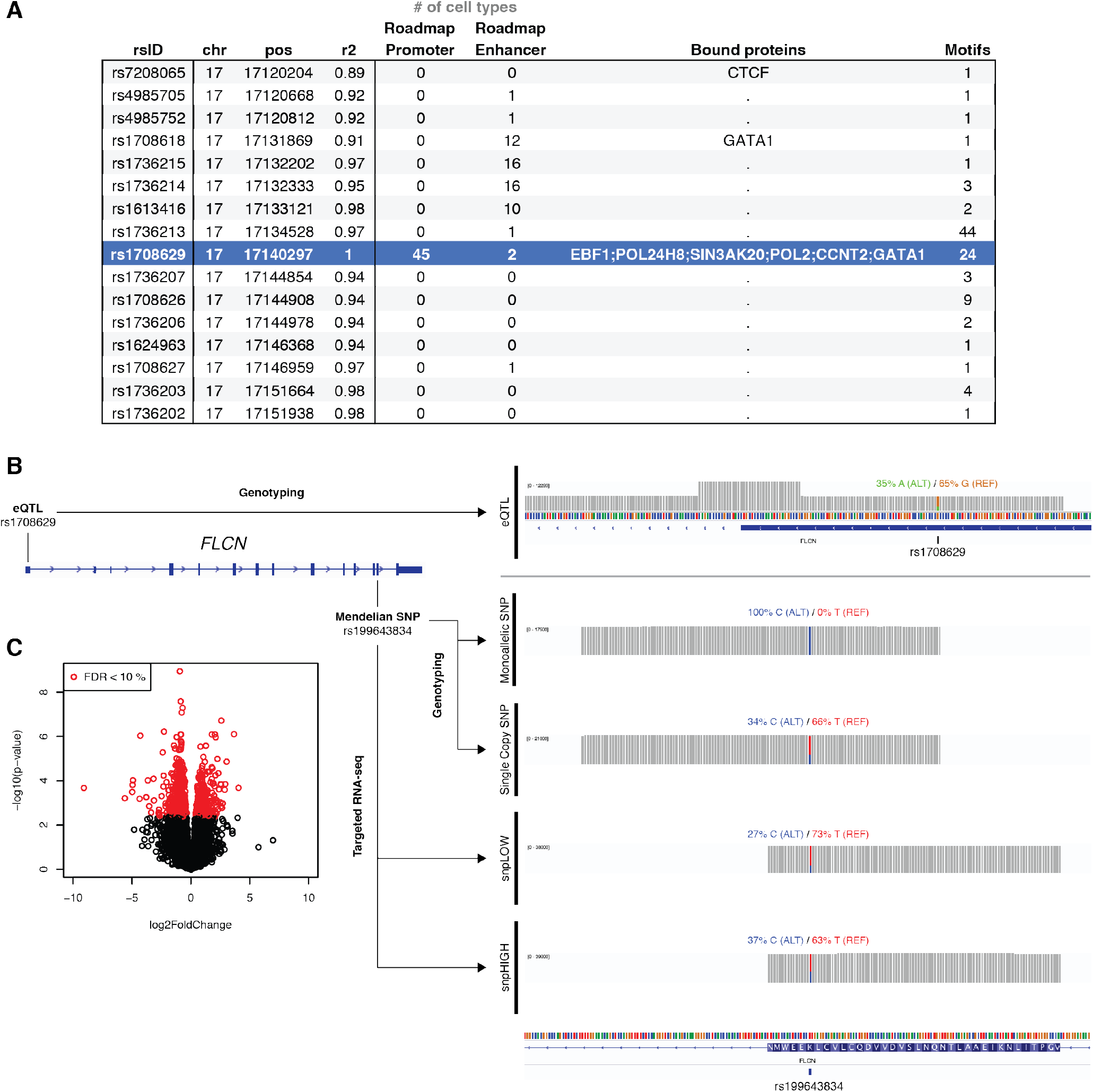
eQTL haplotype aware editing of a Mendelian SNP in 293T cells using the transcriptome as a phenotypic readout. **A)** Haploreg v3 annotations of top causal variant candidates for the *FLCN* eQTL. Highlighted in blue is the variant rs1708629 that we consider as the most likely causal variant, since it is annotated as promoter and enhancer in multiple Roadmap Epigenomic cell types, is bound to numerous proteins, and contains many protein binding motifs. The r2 between the highlighted variant and other variants is listed. **B)** Visualization of eQTL SNP (rs1708629) genotyping reads, edited SNP (rs199643834) genotyping reads from representative monoallelic and single copy clones, and targeted RNA-seq reads from representative low SNP expression (snpLOW) and high SNP expression (snpHIGH) clones. Ratios of the reference allele and alternative allele in targeted DNA and RNA sequencing are indicated. A full list of primers used for sequencing can be found in Table S5. C) Volcano plot of differential expression analysis comparing two clones monoallelic for the edit SNP versus four wildtype clones.

## References

1000 Genomes Project Consortium, Auton, A., Brooks, L.D., Durbin, R.M., Garrison, E.P., Kang, H.M., Korbel, J.O., Marchini, J.L., McCarthy, S., McVean, G.A., et al. (2015). A global reference for human genetic variation. Nature 526, 68–74.

Abecasis, G.R., Altshuler, D.M., Bentley, D.R., Chakravarti, A., Donnelly, P., Eichler, E.E., Green, E.D., Knoppers, B.M., Korbel, J.O., Lander, E.S., et al. (2015). A global reference for human genetic variation. Nature 526, 68–74.

Alberobello, A.T., Congedo, V., Liu, H., Cochran, C., Skarulis, M.C., Forrest, D., and Celi, F.S. (2011). An intronic SNP in the thyroid hormone receptor β gene is associated with pituitary cell-specific overexpression of a mutant thyroid hormone receptor β2 (R338W) in the index case of pituitary-selective resistance to thyroid hormone. Journal of Translational Medicine 2011 9:1–9, 144.

Amin, A.S., Giudicessi, J.R., Tijsen, A.J., Spanjaart, A.M., Reckman, Y.J., Klemens, C.A., Tanck, M.W., Kapplinger, J.D., Hofman, N., Sinner, M.F., et al. (2012). Variants in the 3’ untranslated region of the KCNQ1-encoded Kv7.1 potassium channel modify disease severity in patients with type 1 long QT syndrome in an allele-specific manner. Eur. Heart J. 33, 714–723.

Bolger, A.M., Lohse, M., and Usadel, B. (2014). Trimmomatic: a flexible trimmer for Illumina sequence data. Bioinformatics 30, 2114–2120.

Butt, C., Zheng, H., Randell, E., Robb, D., Parfrey, P., and Xie, Y.-G. (2003). Combined carrier status of prothrombin 20210A and factor XIII-A Leu34 alleles as a strong risk factor for myocardial infarction: evidence of a gene-gene interaction. Blood 101, 3037–3041.

Cancer Genome Atlas Research Network, Weinstein, J.N., Collisson, E.A., Mills, G.B., Shaw, K.R.M., Ozenberger, B.A., Ellrott, K., Shmulevich, I., Sander, C., Stuart, J.M., et al. (2013). The Cancer Genome Atlas Pan-Cancer analysis project. Nat. Genet. 45, 1113–1120.

Castel, S.E., Levy-Moonshine, A., Mohammadi, P., Banks, E., and Lappalainen, T. (2015). Tools and best practices for data processing in allelic expression analysis. Genome Biol. 16, 195.

Castel, S.E., Mohammadi, P., Chung, W.K., Shen, Y., and Lappalainen, T. (2016). Rare variant phasing and haplotypic expression from RNA sequencing with phASER. Nat Commun 7, 12817.

Chen, R., Shi, L., Hakenberg, J., Naughton, B., Sklar, P., Zhang, J., Zhou, H., Tian, L., Prakash, O., Lemire, M., et al. (2016). Analysis of 589,306 genomes identifies individuals resilient to severe Mendelian childhood diseases. Nat. Biotechnol. 34, 531–538.

Cooper, D.N., Krawczak, M., Polychronakos, C., Tyler-Smith, C., and Kehrer-Sawatzki, H. (2013). Where genotype is not predictive of phenotype: towards an understanding of the molecular basis of reduced penetrance in human inherited disease. Hum. Genet. 132, 1077–1130.

Das, S., Forer, L., Schönherr, S., Sidore, C., Locke, A.E., Kwong, A., Vrieze, S.I., Chew, E.Y., Levy, S., McGue, M., et al. (2016). Next-generation genotype imputation service and methods. Nat. Genet. 48, 1284–1287.

Delaneau, O., Howie, B., Cox, A.J., Zagury, J.-F., and Marchini, J. (2013). Haplotype estimation using sequencing reads. Am. J. Hum. Genet. 93, 687–696.

Dimas, A.S., Stranger, B.E., Beazley, C., Finn, R.D., Ingle, C.E., Forrest, M.S., Ritchie, M.E., Deloukas, P., Tavar, S., and Dermitzakis, E.T. (2008). Modifier Effects between Regulatory and Protein-Coding Variation. PLoS Genet. 4, e1000244–10.

Dobin, A., Davis, C.A., Schlesinger, F., Drenkow, J., Zaleski, C., Jha, S., Batut, P., Chaisson, M., and Gingeras, T.R. (2013). STAR: ultrafast universal RNA-seq aligner. Bioinformatics 29, 15–21.

Durbin, R. (2014). Efficient haplotype matching and storage using the positional Burrows-Wheeler transform (PBWT). Bioinformatics 30, 1266–1272.

Edmonson, M.N., Zhang, J., Yan, C., Finney, R.P., Meerzaman, D.M., and Buetow, K.H. (2011). Bambino: a variant detector and alignment viewer for next-generation sequencing data in the SAM/BAM format. Bioinformatics 27, 865–866.

Emison, E.S., McCallion, A.S., Kashuk, C.S., Bush, R.T., Grice, E., Lin, S., Portnoy, M.E., Cutler, D.J., Green, E.D., and Chakravarti, A. (2005). A common sex-dependent mutation in a RET enhancer underlies Hirschsprung disease risk. Nature 434, 857–863.

Fischbach, G.D., and Lord, C. (2010). The Simons Simplex Collection: a resource for identification of autism genetic risk factors. Neuron 68, 192–195.

GTEx Consortium (2013). The Genotype-Tissue Expression (GTEx) project. Nat. Genet. 45, 580–585.

GTEx Consortium (2017). Genetic effects on gene expression across human tissues. Nature 550, 204–213.

Heigwer, F., Kerr, G., and Boutros, M. (2014). E-CRISP: fast CRISPR target site identification. Nat. Methods 11, 122–123.

Iossifov, I., O’Roak, B.J., Sanders, S.J., Ronemus, M., Krumm, N., Levy, D., Stessman, H.A., Witherspoon, K.T., Vives, L., Patterson, K.E., et al. (2014). The contribution of de novo coding mutations to autism spectrum disorder. Nature 515, 216–221.

Irimia, M., Weatheritt, R.J., Ellis, J.D., Parikshak, N.N., Gonatopoulos-Pournatzis, T., Babor, M., Quesnel-Vallières, M., Tapial, J., Raj, B., O’Hanlon, D., et al. (2014). A highly conserved program of neuronal microexons is misregulated in autistic brains. Cell 159, 1511–1523.

Khoo, S.K., Giraud, S., Kahnoski, K., Chen, J., Motorna, O., Nickolov, R., Binet, O., Lambert, D., Friedel, J., Lévy, R., et al. (2002). Clinical and genetic studies of Birt-Hogg-Dubé syndrome. J. Med. Genet. 39, 906–912.

Kircher, M., Witten, D.M., Jain, P., O’Roak, B.J., Cooper, G.M., and Shendure, J. (2014). A general framework for estimating the relative pathogenicity of human genetic variants. Nat. Genet. 46, 310–315.

Krumm, N., Turner, T.N., Baker, C., Vives, L., Mohajeri, K., Witherspoon, K., Raja, A., Coe, B.P., Stessman, H.A., He, Z.-X., et al. (2015). Excess of rare, inherited truncating mutations in autism. Nat. Genet. 47, 582–588.

Lappalainen, T., Montgomery, S.B., Nica, A.C., and Dermitzakis, E.T. (2011). Epistatic selection between coding and regulatory variation in human evolution and disease. Am. J. Hum. Genet. 89, 459–463.

Lek, M., Karczewski, K.J., Minikel, E.V., Samocha, K.E., Banks, E., Fennell, T., O’Donnell-Luria, A.H., Ware, J.S., Hill, A.J., Cummings, B.B., et al. (2016). Analysis of protein-coding genetic variation in 60,706 humans. Nature 536, 285–291.

Liao, Y., Smyth, G.K., and Shi, W. (2014). featureCounts: an efficient general purpose program for assigning sequence reads to genomic features. Bioinformatics 30, 923–930.

Lin, Y.-C., Boone, M., Meuris, L., Lemmens, I., Van Roy, N., Soete, A., Reumers, J., Moisse, M., Plaisance, S., Drmanac, R., et al. (2014). Genome dynamics of the human embryonic kidney 293 lineage in response to cell biology manipulations. Nat Commun 5, 4767.

Loh, P.-R., Danecek, P., Palamara, P.F., Fuchsberger, C., Reshef, Y.A., Finucane, H.K., Schoenherr, S., Forer, L., McCarthy, S., Abecasis, G.R., et al. (2016). Reference-based phasing using the Haplotype Reference Consortium panel. Nat. Genet. 48, 1–8.

Love, M.I., Huber, W., and Anders, S. (2014). Moderated estimation of fold change and dispersion for RNA-seq data with DESeq2. Genome Biol. 15, 550.

McCarthy, S., Das, S., Kretzschmar, W., Delaneau, O., Wood, A.R., Teumer, A., Kang, H.M., Fuchsberger, C., Danecek, P., Sharp, K., et al. (2016). A reference panel of 64,976 haplotypes for genotype imputation. Nat. Genet. 48, 1279–1283.

Milne, R.L., and Antoniou, A.C. (2011). Genetic modifiers of cancer risk for BRCA1 and BRCA2 mutation carriers. Ann. Oncol. 22 Suppl 1, i11–i17.

Mohammadi, P., Castel, S.E., Brown, A.A., and Lappalainen, T. (2017). Quantifying the regulatory effect size of cis-acting genetic variation using allelic fold change. Genome Res. 1–14.

O’Connell, J., Gurdasani, D., Delaneau, O., Pirastu, N., Ulivi, S., Cocca, M., Traglia, M., Huang, J., Huffman, J.E., Rudan, I., et al. (2014). A general approach for haplotype phasing across the full spectrum of relatedness. PLoS Genet. 10, e1004234.

Payne, S.R., and Kemp, C.J. (2005). Tumor suppressor genetics. Carcinogenesis 26, 2031–2045.

Piñero, J., Bravo, À., Queralt-Rosinach, N., Gutiérrez-Sacristán, A., Deu-Pons, J., Centeno, E., García-García, J., Sanz, F., and Furlong, L.I. (2017). DisGeNET: a comprehensive platform integrating information on human disease-associated genes and variants. Nucl Acids Res 45, D833–D839.

Samocha, K.E., Robinson, E.B., Sanders, S.J., Stevens, C., Sabo, A., McGrath, L.M., Kosmicki, J.A., Rehnström, K., Mallick, S., Kirby, A., et al. (2014). A framework for the interpretation of de novo mutation in human disease. Nat. Genet. 46, 944–950.

Sanders, S.J., He, X., Willsey, A.J., Ercan-Sencicek, A.G., Samocha, K.E., Cicek, A.E., Murtha, M.T., Bal, V.H., Bishop, S.L., Dong, S., et al. (2015). Insights into Autism Spectrum Disorder Genomic Architecture and Biology from 71 Risk Loci. Neuron 87, 1215–1233.

Sanjana, N.E., Shalem, O., and Zhang, F. (2014). Improved vectors and genome-wide libraries for CRISPR screening. Nat. Methods 11, 783–784.

Snozek, C.L.H., Lagerstedt, S.A., Khoo, T.K., Rubenfire, M., Isley, W.L., Train, L.J., and Baudhuin, L.M. (2009). LDLR promoter variant and exon 14 mutation on the same chromosome are associated with an unusually severe FH phenotype and treatment resistance. Eur. J. Hum. Genet. 17, 85–90.

Toro, J.R., Wei, M.-H., Glenn, G.M., and Weinreich, M. (2008). BHD mutations, clinical and molecular genetic investigations of Birt–Hogg–Dubé syndrome: a new series of 50 families and a review of published reports. J Med Genet 45, 321–331.

Wang, J., Vasaikar, S., Shi, Z., Greer, M., and Zhang, B. (2017). WebGestalt 2017: a more comprehensive, powerful, flexible and interactive gene set enrichment analysis toolkit. Nucl Acids Res 45, W130–W137.

Ward, L.D., and Kellis, M. (2012). HaploReg: a resource for exploring chromatin states, conservation, and regulatory motif alterations within sets of genetically linked variants. Nucl Acids Res 40, D930–D934.

Wei, W.-H., Hemani, G., and Haley, C.S. (2014). Detecting epistasis in human complex traits. Nat. Rev. Genet. 15, 722–733.

Werling, D.M., Brand, H., An, J.-Y., Stone, M.R., Glessner, J.T., Zhu, L., Collins, R.L., Dong, S., Layer, R.M., Markenscoff-Papadimitriou, E.-C., et al. (2017). Limited contribution of rare, noncoding variation to autism spectrum disorder from sequencing of 2,076 genomes in quartet families. bioRxiv 127043.

Zhao, M., Kim, P., Mitra, R., Zhao, J., and Zhao, Z. (2015). TSGene 2.0: an updated literature-based knowledgebase for tumor suppressor genes. Nucl Acids Res 44, D1023–D1031.

